# Quercetin and Fisetin activate circadian clock via RORα and inhibit adipocyte growth

**DOI:** 10.64898/2026.04.28.721484

**Authors:** Xuekai Xiong, Jemima Pangemanan, Tali Kiperman, Zuoming Sun, Antoni Paul, Vijay Yechoor, Ke Ma

## Abstract

The circadian clock maintains temporal control of metabolic processes and exerts a key role in adipocyte development. Discovery of clock-modulatory compounds may provide new avenues for metabolic disease therapy. Here we report the identification of flavonoid compounds, Quercetin and Fisetin, as clock-activating molecules with direct inhibitory action on adipogenesis and adipocyte lipid metabolism. Quercetin and Fisetin displayed robust RORα agonism that promoted clock oscillation with induction of clock genes. Treating preadipocytes with these compounds blocked their adipogenic differentiation. In mature adipocytes, Quercetin and Fisetin suppressed lipid accumulation by inhibiting lipogenic enzymes. Furthermore, activation of RORα by a synthetic agonist or ectopic expression were sufficient to inhibit adipogenesis. In mice treated with Quercetin or Fisetin, RORα was markedly induced in adipose depots with strong suppression of the adipogenic and lipogenic programs. While quercetin significantly attenuated lipid storage in adipose tissue in vivo accompanied with lowering of free fatty acids and improved insulin sensitivity, fisetin displayed a less robust effect with differential regulation of lipolytic pathway. Collectively, these findings uncovered the clock-activating properties of quercetin and fisetin that prevent adipocyte maturation and hypertrophy to limit adipose tissue expansion. These actions contribute, at least in part, to their beneficial effects on metabolic disorders.

## Introduction

The cell-autonomous circadian clock drives ∼24 hour oscillations of key metabolic pathways that are essential to maintain homeostasis by providing appropriately-timed responses to nutrient cues [1, 2]. Disruption of the clock control or the timing cues predispose to metabolic dysregulations, and is believed to be a significant contributing factor to the current epidemic of obesity and Type II diabetes [3–5]. Clock modulation of metabolic processes and its role in the etiology of metabolic disorders suggests it as a potential therapeutic target [6–8]. Identification of clock-modulatory molecules, particularly targeted interventions toward re-enforcing the metabolic beneficial effects of circadian regulation, could lead to novel therapies for obesity or anti-diabetes drugs [9–12].

The circadian clock rhythm is generated by a transcriptional/translational feedback circuit composed of positive and negative regulatory components [1]. Transcription activators CLOCK (Circadian Locomotor Output Cycles Kaput) and Bmal1 (Brain and Arnt-Like 1) initiates clock transcription that is countered by feedback inhibition by their direct target genes that are transcription repressors, Periods (Per 1-3) and Cryptochromes (Cry1 & 2), which interact with Bmal1/CLOCK to repress transcription. In addition, RORα (Retinoid-related Orphan Receptor α) and Rev-erbα, a pair of antagonistic transcription factors, exert positive and negative regulations, respectively, of Bmal1 rhythmic expression and thus constitutes a re-enforcing loop to ensure robustness of clock oscillation [13, 14]. Interestingly, both the positive and negative arms of the core molecular clock machinery are involved in modulating specific stages of adipocyte development. *CLOCK* mutant mice [15] or loss of the transcription activator *Bmal1* resulted in the development of obesity [16, 17], impacting adipogenic development or fatty acid metabolism. On the other hand, a key component of the negative arm of the clock feedback loop, PER2, can interfere with the adipogenic activity of PPARγ to block block adipocyte differentiation that impacts lipid metabolism [18].

Through orchestration of time-of-day dependence of key metabolic pathways in distinct metabolic organs, circadian temporal control is intimately linked with metabolic homeostasis [2]. Circadian misalignment predisposes to obesity and insulin resistance, as established by large-scale epidemiological investigations and accumulating experimental evidence [4, 19–23]. Significant dampening of clock oscillation amplitude occurs with nutritional overload in high-fat diet-induced obesity, and clock disruption could be synergistic in inducing diabetes with over-nutrition [24, 25]. It is thus conceivable that pharmacological targeting of key clock regulators to maintain or re-enforce clock-controlled metabolic rhythms may provide new avenues for anti-obesity therapies. Recent efforts in identifying molecules with clock-modulatory activities indeed demonstrated their potential for applications in metabolic diseases [9–12]. As readily “druggable” nuclear receptors within the molecular clock regulatory circuit, the re-enforcing loop driven by the reciprocal positive and negative regulatory control through RORα and Rev-erbα, respectively, has been explored [9, 26, 27]. Notably, both agonists and antagonists of the clock repressor Rev-erbα, a direct negative transcriptional target of CLOCK/Bmal1, were identified [28]. Agonism of Rev-erbα by synthetic ligands reveled their therapeutic efficacy in protecting against obesity in mice that ameliorated dyslipidemia [29]. Nobiletin, a citrus flavonoid identified as a ROR-activating natural ligand [11], exhibit direct inhibitory effect on adipogenesis with demonstrated anti-obesity efficacy in vivo [30]. Thus, targeting the circadian clock function involved in adipocyte development may lead to the development of novel pharmacological interventions for obesity and related metabolic consequences.

Quercetin and Fisetin belong to a class of flavonoid compounds that are highly abundant in fruit sources [31]. These natural compounds are associated with numerous metabolic benefits and have been shown to reduce the risk for cardiovascular disease, dyslipidemia and Type II diabetes [32–34]. Notably, Quercetin and Fisetin are known to possess anti-aging effects through selective induction of apoptosis in senescent cells [32], leading to senolytic activity with epidemiolocal evidence supporting the association with their anti-aging properties [35]. The combination of Quercetin and Dasatinib has shown strong efficacy in reducing senescent cell burden and is currently being tested in clinical trials for various anti-aging effects [36–39], though a multitude of additional beneficial mechanisms involving the anti-inflammatory and anti-oxidant actions of Quercetin may also apply [40]. Interestingly, our recent study revealed that Nobiletin, a structurally related flavonoid compound with clock amplitude-enhancing properties via RORα/γ activation [11], exerts direct suppression of adipogenic progenitor maturation with strong effects on reducing adipose tissue mass in vivo [30].

Based on the shared flavonoid structure between Quercetin and Fisetin with Nobiletin, we explored their potential clock modulatory actions and effects on adipocyte maturation and lipid metabolism. Through investigation of the activities of Quercetin and Fisetin using cellular models coupled with in vivo validation, the current study uncovered their RORα agonist function and augmented clock oscillation with inhibition of adipocyte development and hypertrophy.

## Materials & Methods

### Animal studies

Mice were maintained in the City of Hope vivarium under a constant 12:12 light dark cycle and were provided with nestlets and maze for cage enrichment. All animal experiments were approved by the Institutional Animal Care & Use Committee (IACUC) of City of Hope and performed according to the IACUC approval. Eight-week-old C57BL/6J mice purchased from Jackson Laboratory were used following 2 weeks of acclimation. Quercetin (CAS 849061-97-8) and fisetin (CAS 528-48-3) were purchased from Cayman Chemical Santa Cruz (CAS 480-41-1). They were administered to mice at 50 mg/kg dose via daily intraperitoneal injections for 5 days.

### Cell culture

3T3-L1 preadipocytes (RRID:CVCL 0123) were obtained from ATCC, and maintained in DMEM with 10% fetal bovine serum supplemented with 1% Penicillin-Streptomycin-Glutamine, as previously described (26, 27). Cells were seeded at 1×10^6^ density on 6-well plates and treated with quercetin or fisetin (5uM and 10μM) or vehicle (DMSO) for 6 hours prior to RNA extraction.

### Lentivirus packaging and infection for generation of stable cell line

Full-length *Rora* was constructed by subcloning the corresponding cDNA into pcDNA3.0-Flag or pCDH-puro vector. The 293A human embryonic kidney cells were transfected with the packaging plasmids (pSPAX.2 and pMD2.G) and recombinant lentivirus vectors using the PEI reagent according to the manufacturer’s protocol. After 48 h post-transfection, lentiviruses were collected through 0.45 μm filter to remove the cell debris. The 3T3-L1 preadipocytes were infected by the lentivirus medium supplemented with 8 μg/mL polybrene. After the 24 h infection, stable cell lines were selected in the presence of 2 μg/mL puromycin for selection of stable clones. The RORα overexpressing stable cell line and its vector control were verified by RT-qPCR and immunoblotting.

### Adipogenic differentiation

Normal 3T3-L1, pCDH control and Rorα-overexpressing stable 3T3-L1 cells were subjected to differentiation in 6-well plates at 90% confluency. For adipogenic differentiation, induction media containing 1.6μM insulin, 1μM dexamethasone, 0.5mM IBMX and 0.5 uM Rosiglitazone was used for 3 days followed by maintenance medium with insulin for 3 days for 3T3-L1, as previously described [30, 41]. SR1078 (CAS 1246525-60-9) and SR3335 (CAS 293753-05-6) were purchased from Cayman Chemical. For treatments with SR1078 and SR3335 (2 µM) or vehicle (DMSO) of 3T3-L1 preadipocytes, these compounds were administered for the entire differentiation time course prior to staining and protein extraction.

### Primary preadipocyte isolation and treatment

The stromal vascular fraction containing preadipocytes were isolated from subcutaneous fat pads, as previously described [42]. Briefly, fat pads were cut into small pieces and digested using 0.1% collagenase Type 1 with 0.8% BSA at 37^0^C with constant shaking for 60 minutes. The digested homogenate was passed through Nylon mesh and centrifuged to collect the pellet containing the stromal vascular fraction with preadipocytes. Pelleted preadipocytes were cultured in F12/DMEM supplemented with bFGF (2.5 ng/ml), expanded for two passages and subjected to differentiation in 6-well plates at 90% confluency. Primary preadipocytes were treated with quercetin and fisetin (5uM and 10μM) or vehicle (DMSO) for 6 hours prior to RNA extraction or 24 hours prior to protein extraction.

### Primary preadipocyte differentiation

Primary preadipocytes obtained from the stromal vascular fraction of mice subcutaneous fat pads were subjected to differentiation in 6-well plates at 90% confluency. Adipogenic differentiation was induced for 2 days in medium containing 10% FBS, 1.6 μM insulin, 1 μM dexamethasone, 0.5 mM IBMX, 0.5 uM rosiglitazone before switching to maintenance medium for 4 days with insulin only. Compounds at indicated concentrations were administered for the entire differentiation time course following adipogenic induction or added at 4 day differentiated adipocytes for 2 days prior to staining and protein extraction.

### Oil-red-O and Bodipy staining

Staining of neutral lipids following adipogenic differentiation at indicated times were performed as previously described [43]. Briefly, for oil-red-O staining, cells were fixed with 10% formalin and incubated in 0.5% oil-red-O solution for 1 hour. Bodipy 493/503 was used at 1mg/L together with DAPI for 15 minutes, following 4% paraformaldehyde fixation and permeabilization with 0.2% triton-X100.

### Continuous Bioluminescence monitoring of Per2::dLuc luciferase reporter

3T3-L1, C3H10T1/2 or primary preadipocytes containing a *Per2::dLuc* luciferase reporter were used for bioluminescence recording, as previously described [30]. Cells were seeded at 4×10^5^ density on 24 well plates at 90% confluence following overnight culture with explant medium luciferase recording media. Explant medium contains DMEM buffer stock, 10% FBS, 1% PSG, pH7 1M HEPES, 7.5% Sodium Bicarbonate, Sodium Hydroxide (100mM) and XenoLight D-Luciferin bioluminescent substrate (100mM). Raw and subtracted results of real-time bioluminescence recording data for 6 days were exported, and data was calculated as luminescence counts per second, as previously described [44]. LumiCycle Analysis Program (Actimetrics) was used to determine clock oscillation period, length amplitude and phase. Briefly, raw data following the first cycle from day 2 to day 5 were fitted to a linear baseline, and the baseline-subtracted data (polynomial number = 1) were fitted to a sine wave, from which period length and goodness of fit and damping constant were determined. For samples that showed persistent rhythms, goodness-of-fit of >80% was usually achieved.

### ROR-responsive luciferase reporter assay

RORE-containing luciferase reporter containing three RORE bindings sites RORE(3) TK-Luc was a gift from Zuoming Sun lab [45, 46]. 293T cells were seeded and grown overnight to ∼70% confluency prior to plasmid transfection. Cells were transfected using PolyJet reagent (SignaGen Laboratories) with expression plasmids, including RORE-luciferase reporter (50 ng/well), Renilla (20 ng/well), and RORα (20 ng/well) following the manufacturer’s protocol. 24 hours following transfection, cells were treated with flavonoids or SR1078 at indicated concentrations overnight to induce luciferase activity. Luciferase activity was assayed using Dual-Luciferase Reporter Assay Kit (Promega) in 96-well black plates. Firefly luciferase reporter luminescence was measured on microplate reader (TECAN infinite M200pro) and normalized to control Renilla luciferase activity, as previously described [17]. The mean and standard deviation values of four repeats were calculated for each well and graphed.

### Hematoxylin and eosin histology

Adipose tissues were fixed in 10% neutral-buffered formalin for 72 hours prior to embedding. 10 μm paraffin sections were processed for hematoxylin and eosin staining.

### Plasma metabolite analysis

Plasma levels of glucose (Thermo Scientific), triglyceride (Teco Diagnostics) and free fatty acids (BioAssay Systems) were measured using 5-10μl of plasma samples with respective commercial kits, according to manufacturer’s protocols.

### Insulin tolerance test

Insulin tolerance test (ITT) was performed in mice fasted briefly for 4 hours, as described previously [47]. 0.5 U/kg insulin was injected intraperitoneally, and glucose levels were measured at indicated times via tail vein bleed using a glucometer (AimStrip Plus).

### Immunoblot analysis

Total protein was extracted using lysis buffer containing 3% NaCl, 5% Tris-HCl, 10% Glycerol, 0.5% Triton X-10 with protease inhibitor cocktail. 20-40 µg of total protein was resolved on 10% SDS-PAGE gels followed by immunoblotting on PVDF membranes (Bio-rad). Membranes were developed by chemiluminescence (SuperSignal West Pico, Pierce Biotechnology) and signals were obtained via a chemiluminescence imager (Amersham Imager 680, GE Biosciences). Primary antibodies used are listed in Supplemental Table 1.

### RNA extraction and RT-qPCR analysis

PureLink RNA Mini Kit (Invitrogen) were used to isolate total RNA from cells. cDNA was generated using Revert Aid RT kit (ThermoFisher) and quantitative PCR was performed using SYBR Green Master Mix (Thermo Fisher) in triplicates on ViiA 7 Real-Time PCR System (Applied Biosystems). Relative gene expression was calculated using the comparative Ct method with normalization to 36B4 as internal control. PCR primer sequences are listed in Supplemental Table 2.

### Statistical analysis

Data are presented as mean ± SD. Each experiment was repeated at minimum three times to validate the result. The number of replicates for each experiment were indicated in figure legends. Two-tailed Student’s t-test or One-way ANOVA with post-hoc analysis for multiple comparisons were performed as appropriate using GraphPad PRISM as indicated. P<0.05 was considered statistically significant.

## Results

### Clock-modulatory activities of Quercetin and Fisetin in adipogenic progenitors via RORα activation

Both quercetin and fisetin belong to the flavonol subclass of flavonoid compounds with a hydroxyl group at C-3 of the central ring, as well as additional hydroxyl groups in rings A and B that distinguish them from the polymethoxyflavone Nobiletin (Fig. 1A). The only structural difference between quercetin and fisetin is a hydroxyl group at position 5 of the A ring present in quercetin, but not in fisetin, which may render fisetin less polar (Fig. 1A). Nobiletin is known to activate RORα that enhances clock cycling amplitude and displays anti-obesity effect in vivo [11, 30]. Based on the shared common flavonoid structure with nobiletin, we determined whether quercetin and fisetin can modulate RORα and clock activity. The effect of quercetin and fisetin on clock modulation was determined using continuous real-time luciferase monitoring of preadipocytes containing a *Per2::dLuc* reporter isolated from transgenic mice. Notably, Quercetin treatment at 5 and 10 µM led to dose-dependent increases in clock cycling amplitude (Fig. 1B) coupled with reductions of period length, indicating activation of clock oscillation. In 3T3-L1 preadipocytes, quercetin was able to induce Bmal1 and CLOCK protein levels (Fig. 1C), further supporting its activation of clock in adipogenic precursors. Fisetin also displayed clock-activating effects in *Per2::dLuc* preadipocytes, with dose-dependent inductions of clock amplitude together with period shortening (Fig. 1D). Interestingly, as compared with quercetin, the effect of fisetin at high concentrations appeared to reach a plateau at between 5 to 10 µM. Analysis of clock proteins also revealed inductions of Bmal1 and CLOCK protein by Fisetin, albeit only at 10 µM (Fig. 1E). We next determined the effects of quercetin and fisetin functions on RORα using a RORE-containing luciferase reporter (Kane & Means, 2000), together with nobiletin and a synthetic RORα agonist SR1078 as positive controls (Wang et al., 2010). RORE-driven luciferase reporter (Fig. 1F). As expected, ectopic expression of RORα induced up to ∼18-fold induction of its activity over empty vector control (Fig. 1F). As anticipated, the synthetic RORα agonist SR1078 induced activation of the luciferase reporter, while Nobiletin displayed lower potency at compared to SR1078. Quercetin and fisetin at 2µM were both able to induce RORE-driven luciferase to a similar degree as 1 µM of SR1078, indicating they indeed have RORα agonist activity (Fig. 1F). Importantly, although their activities were less potent than SR1078, at 2µM both quercetin and fisetin displayed a higher luciferase induction than that of nobiletin. Interestingly, dose-response assessment of quercetin and fisetin activation of RORE-luc reporter revealed that their effects reached maximal at 2 µM without further activation at higher concentrations, and fisetin appeared to lose its agonist activity at 5 and 10 µM (Fig. 1G).

**Figure 1.**
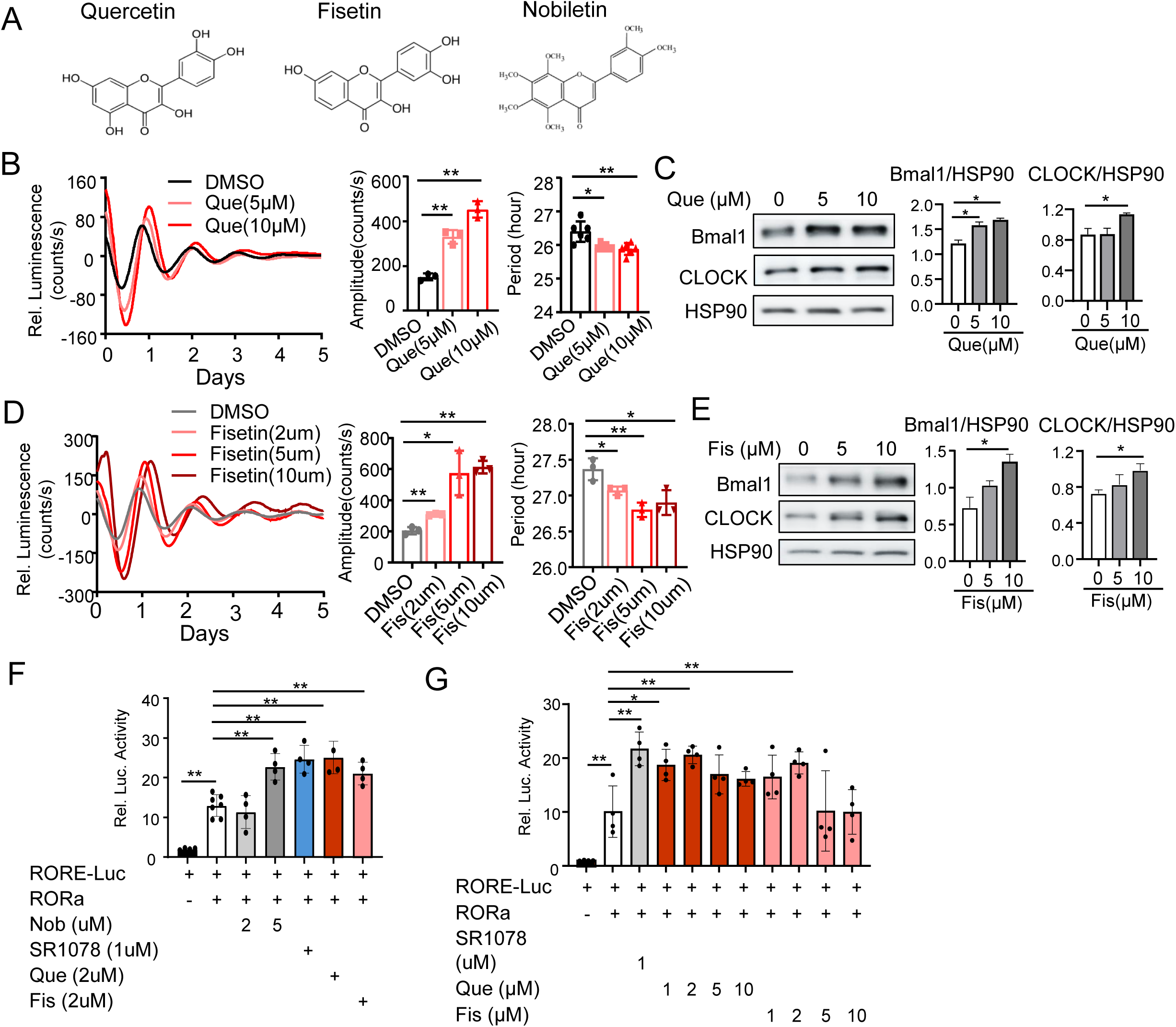
Clock-modulatory activities of Quercetin and Fisetin in adipogenic precursors via activation of RORα. (A) Chemical structure of Quercetin and Fisetin, with comparison to Nobiletin. (B, D) Effect of Quercetin (B) and Fisetin (D) on clock modulation, with baseline-adjusted tracing plots of average luciferase bioluminescence activity and quantitative analysis of oscillatory amplitude and period length. Luciferase activity was monitored for 6 days using preadipocytes containing Per2-dLuc reporter (n=3). *, **: p<0.05 and 0.01 vs. DMSO by Student’s t test. (C, E) Immunoblot analysis of core clock protein induction by Quercetin (C) and Fisetin (E) in primary preadipocytes following 16 hours of treatment at indicated concentrations, with corresponding quantitative analyses. Data are presented as Mean ± SD of n=3 replicates. (F, G) Effect of Quercetin and Fisetin on RORE-driven luciferase reporter activity, as compared with nobiletin or a synthetic ligand SR1078 (F), and dose-response analysis of these compounds (G). Data are presented as Mean ± SD of n=4 replicates. *, **: p<0.05 or 0.01 by Student’s t test.

### Quercetin and Fisetin suppress adipogenesis

Findings of quercetin and fisetin activities as RORα agonists led us to further examine their effects on transcriptional regulation of clock genes. In 3T3-L1 preadipocytes, acute treatment by quercetin or fisetin for 6 hours revealed significant comparable inductions of RORα, but not other core clock regulators Bmal1 and CLOCK (Fig. 2A). This transcriptional activation of RORα by quercetin and fisetin suggests a potential positive regulatory loop via their agonist activity. The inductions of RORα by these compounds were further demonstrated by protein analysis using 16-hour treated samples (Fig. 2B & 2C), along with elevated CLOCK or Bmal1 protein levels. Furthermore, in primary preadipocytes isolated from adipose tissue stromal vascular fraction, we found similar effects of quercetin and fisetin on inducing RORα transcript and protein expression (Fig. 2D). Based on the inhibition of clock on adipogenesis and that the RORα-activating nobiletin displayed clock-dependent effect on adipogenic differentiation [30], we determined potential direct modulation of adipogenesis by quercetin and fisetin via treatment using primary preadipocytes undergoing adipgenic differentiation. As shown by oil-red-O and Bodipy staining to assess lipid accumulation as a definitive marker for mature adipocyte formation, both compounds markedly inhibited the adipogenic maturation of preadipocytes subjected to a standard differentiation time course of 6 days (Fig. 2E & 2F). Based on the quantitative analysis of Bodipy staining, quercetin displayed stronger inhibition of adipocyte differentiation than that of fisetin. While quercetin was sufficient to exert ∼45% and 70% of adipogenic inhibition at 2 and 10 uM, respectively (Fig. 2E), fisetin achieved only ∼31% and 48% at comparable concentrations (Fig. 2F).

**Figure 2.**
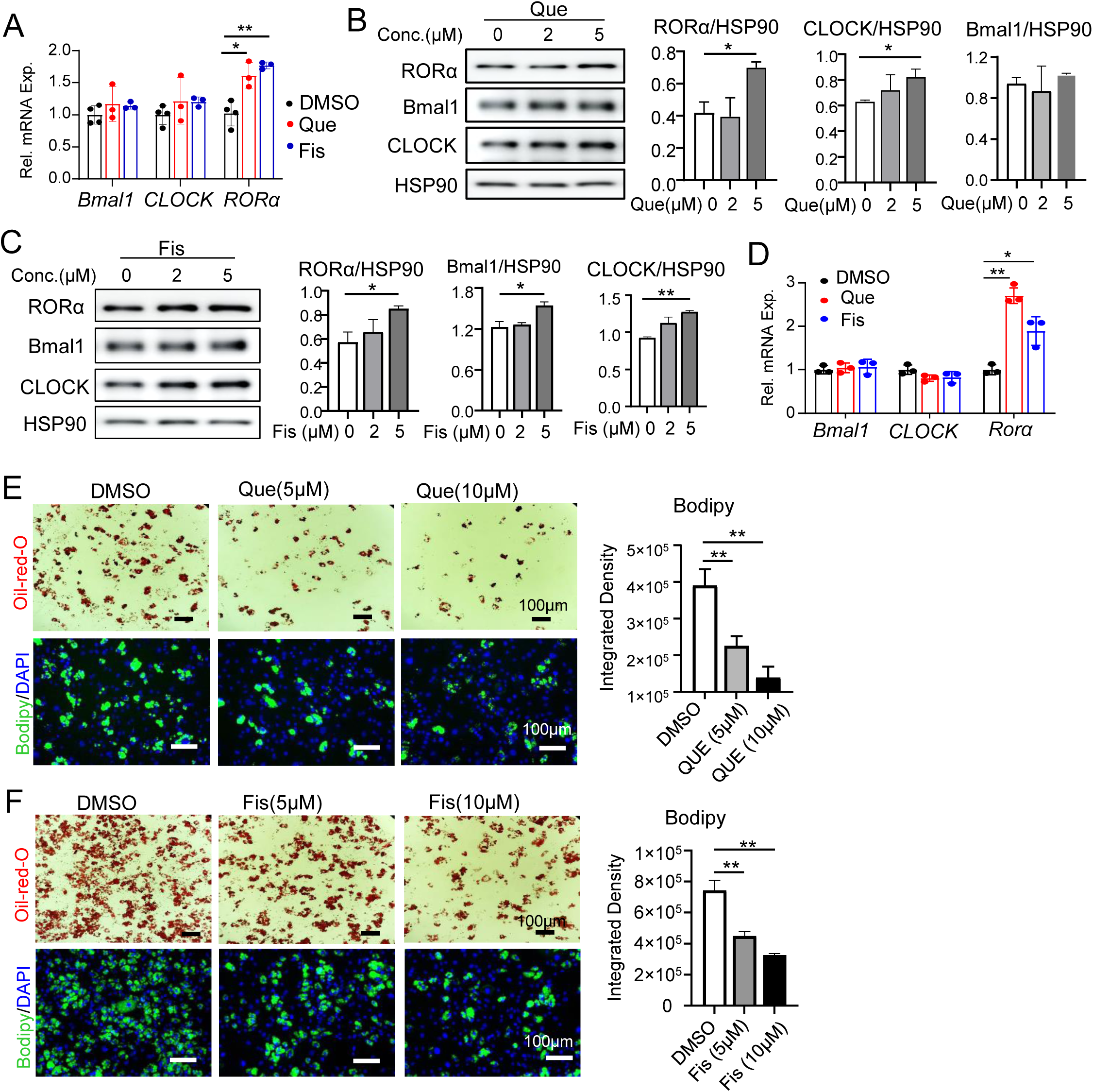
Quercetin and Fisetin inhibition of adipogenic progenitor differentiation. (A) RT-qPCR analysis of Quercetin and Fisetin treatment for 6 hours 5µM on clock gene modulation in 3T3-L1 preadipocytes for 6 hours. (B, C) Representative immunoblot analysis of effect of Quercetin (B) or Fisetin (C) following 16 hours of treatment in 3T3-L1 preadipocytes at indicated concentrations with corresponding quantitative analyses. Data are presented as Mean ± SD of n=3 replicates. (D) RT-qPCR analysis of effect of 6-hour treatment of Quercetin (5µM) or Fisetin (5µM) on clock gene expression in mouse primary preadipocytes isolated from the stromal vascular fraction of inguinal adipose fat pad (n=3/group). (E, F) Representative images of oil-red-O (upper panel) and Bodipy staining (lower panel) of primary mouse preadipocyte treated with Quercetin (E), or Fisetin (F), at indicated concentrations at day 6 of adipogenic differentiation, Quantitative analysis of BODIPY fluorescence of three representative fields were performed with normalization to cell number as shown by DAPI nuclear stain. Scale bars: 100 μm. *, **: p<0.05 and 0.01 vs. DMSO by Student’s t test.

### Quercetin and Fisetin inhibit lipid accumulation in differentiated mature adipocytes

Mature adipocytes compose the predominant portion of adipose tissue mass. In adult adipose depots, adipocyte hypertrophy as a result of excess lipid storage is a major mechanism driving fat mass expansion in obesity. We exposed differentiated adipocytes derived from preadipocytes to different concentrations of quercetin or fisetin and examined lipid dynamics in these mature adipocytes. As shown by Bodipy assessment of the total amount of lipids in differentiated adipocytes, quercetin treatment for 24 hours was able to markedly block lipid accumulations in a dose-dependent manner with ∼60% of total inhibition at 10 µm (Fig. 3A & 3B). Quercetin treatment resulted in a near complete suppression of the lipogenic enzyme fatty acid synthase (FASN) together with marked loss of adipogenic and adipocyte markers (Fig. 3C). Notably, though PPARγ protein displayed significant reductions by treatment with quercetin, C/EBPα was not altered. The effect of Fis on lipid accumulation in adipocytes was comparatively more modest, with strongest inhibition at 10 µM at less than 40% (Fig. 3 D & 3E), while its regulation on lipogenic enzymes were not significant except inhibition of FABP4 (Fig. 3F). Together, these findings revealed the direct suppression of quercetin and fisetin on lipid storage in mature adipocytes, with quercetin demonstrating a stronger efficacy, suggesting that their inhibition of adipocyte hypertrophic growth could contribute to anti-obesity applications [48].

**Figure 3.**
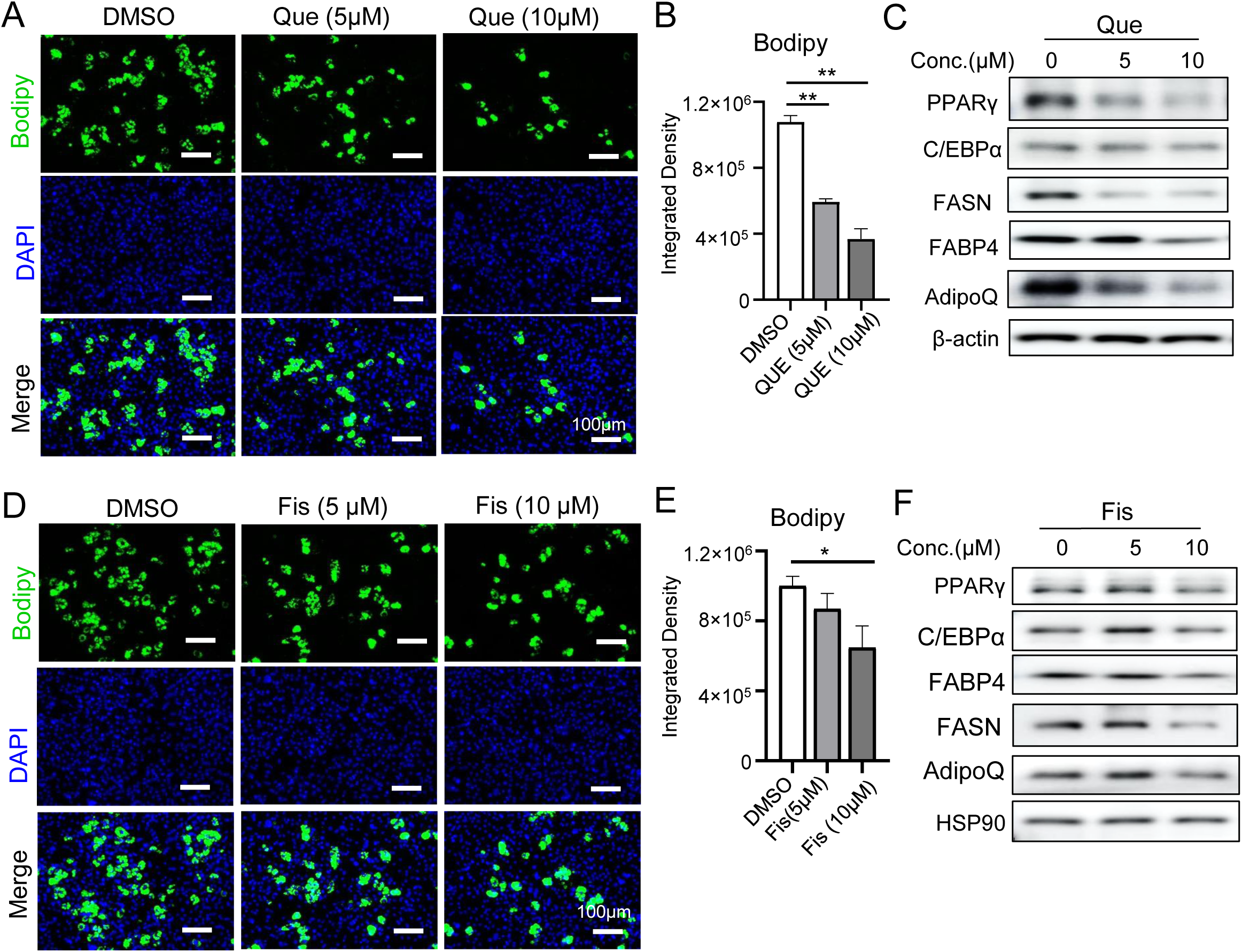
Effect of Quercetin and Fisetin on lipid storage in mature adipocytes. (A, B, D, E) Representative images of Bodipy fluorescence staining of lipids in mature adipocytes treated with Quercetin (A), or Fisetin (D), with quantitative analyses of intensity (B & E). Adipocytes were differentiated from primary preadipocytes for 6 days, followed by Quercetin and Fisetin treatment for 48 hours at indicated concentrations. Corresponding quantitative analysis of staining intensity of Bodipy were shown in (B) and (E), respectively. (C, F). Representative images of immunoblot analysis of audiogenic and lipogenic proteins of mature adipocytes treated by Quercetin (C), or Fisetin (F).

### Pharmacological activation of RORa by SR1078 suppresses adipogenesis

Findings of RORα agonist activities of quercetin and fisetin and their effect on inhibiting adipogenic differentiation led us to postulate that pharmacological interventions of RORα function via synthetic ligands may exert similar effect on adipocyte development. Using *Per2::dLuc* preadipocytes, we first validated that the RORα agonist SR1078 was able to induce clock amplitude (Fig. 4A & 4B). When administered to 3T3-L1 preadipocytes undergoing adipogenic differentiation, SR1078 resulted in nearly complete blockade of lipid accumulation, indicative of inhibition of adipocyte maturation (Fig. 4C). In contrast, pharmacological inhibition of RORα using the antagonist SR3335 was sufficient to promote adipogenesis, as shown by elevated levels of lipid staining by Bodipy (Fig. 4D), which was validation by quantitative analysis.

**Figure 4.**
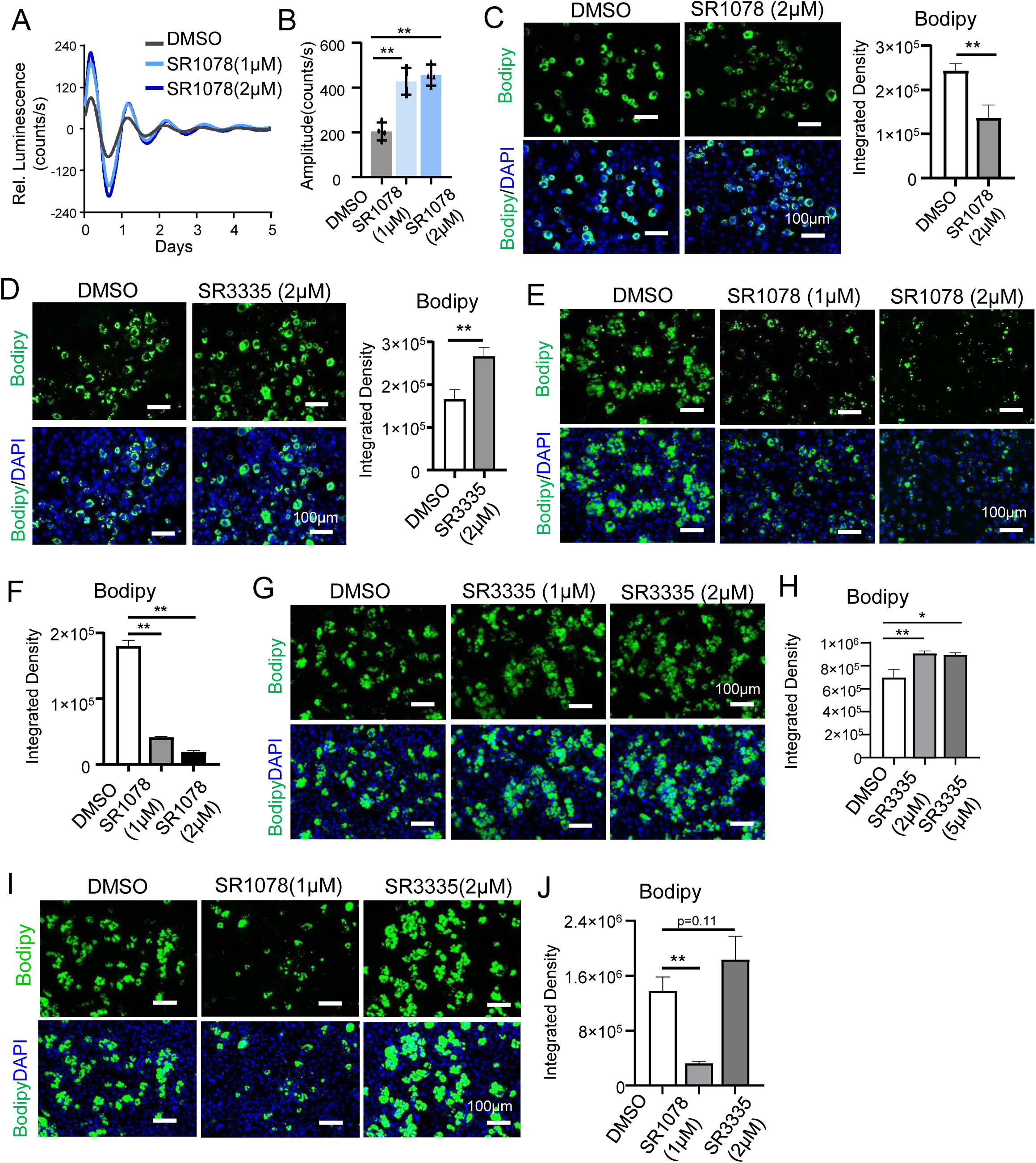
Effect of RORα pharmacological modulation on adipogenesis. (A, B) The effect of synthetic RORα agonist SR1078 treatment on clock modulation in 3T3-L1 preadipocytes, as shown by baseline-adjusted tracing plots of average luciferase bioluminescence activity (A), with quantitative analysis of oscillatory amplitude (B). (C, D) Representative images of Bodipy staining of 3T3-L1 adipocytes treated with SR1078 (C, 2µM), or SR3335 (D, 2µM), at day 6 differentiation with quantitative analysis of Bodipy (n=3/group). (E-H) The effect of SR0178 or SR3335 treatment on differentiation of primary preadipocyte as shown by representative images of Bodipy staining at day 6 (E & G), with quantitative analysis (). Quantitative analysis of Bodipy staining was shown in F & H (n=3/group). (I, J) Representative images of Bodipy staining of day 5-differnetiated primary adipocytes treated with indicated concentrations of SR1078 or SR3335 for 24 hours (I), with corresponding quantitative analyses (J, n=3/group). Scale bar: 100 μm. *, **: p<0.05 or 0.01 vs. DMSO by Student’s t test.

Similar effects of SR1078 on inhibiting adipogenesis were observed in primary preadipocytes undergoing adipogenic differentiation, as shown by both Bodipy staining (Fig. 4E) and quantitative analysis (Fig. 4F). Treatment with 1 or 2 uM of SR1078 resulted in near 80-90% reduction in lipid staining (Fig. 4E & 4F), indicating an almost complete block of adipogenic differentiation. In line with findings from 3T3-L1 adipocytes, treatment of primary preadipocytes with SR3335 led to enhanced differentiation as shown by lipid staining (Fig. 4G & 4H). Similarly as tested for quercetin and fisetin, we next determined whether these synthetic ligands modulate lipid dynamics in mature adipocytes following differentiation. The overall impact of SR1078 and SR3335 on mature adipocytes, as shown in Fig. 4I and 4J, were largely concordant with their distinct modulations of adipogenesis. While SR1078 inhibited lipid accumulation in differentiated adipocytes following treatment for 48 hours of treatment, SR3335 displayed a tendency toward increasing lipid content in mature adipocytes (Fig. 4I & 4J). Thus, these findings from pharmacological activation or inhibition of RORα suggest its coordinated modulation of adipocyte development and hypertrophy, with its activation preventing while the antagonism enhancing adipogenic maturation and lipid storage.

### Genetic RORa gain-of-function blocks adipogenic differentiation

We next further tested whether promoting RORα function in preadipocytes affects adipogenic differentiation via ectopic expression. RORα was overexpressed in 3T3-L1 cells using a flag-tagged cDNA expression vector that resulted in robust expression, as well as inductions of core clock genes *Bmal1* and *Nr1d1* (Fig. 5A). Protein expression of Flag-RORα was confirmed in these stable cell lines, along with induction of endogenous protein levels, suggesting a positive feed forward loop involved in RORα transcription (Fig. 5B). Bmal1 and Nr1d1 proteins were increased similarly as their transcripts, indicating the activation of clock by forced expression of RORα. When subjected to adipogenic differentiation, cells with RORα overexpression displayed a clear lack of mature adipocyte formation as compared to cells with vector controls, as indicated by oil-red-O (Fig. 5C) and Bodipy (Fig. 5D) staining. Consistent with the marked inhibition of adipogenic maturation, analysis of adipogenic gene induction at day 6 of adipocyte differentiation of these cells revealed the lack of FABP4, a mature adipocyte marker as well as loss of adipogenic factors C/EBPα and PPARγ (Fig. 5E). Taken together, these results from genetic gain-of-function studies of RORα further demonstrated its role in inhibiting adipocyte development.

**Figure 5.**
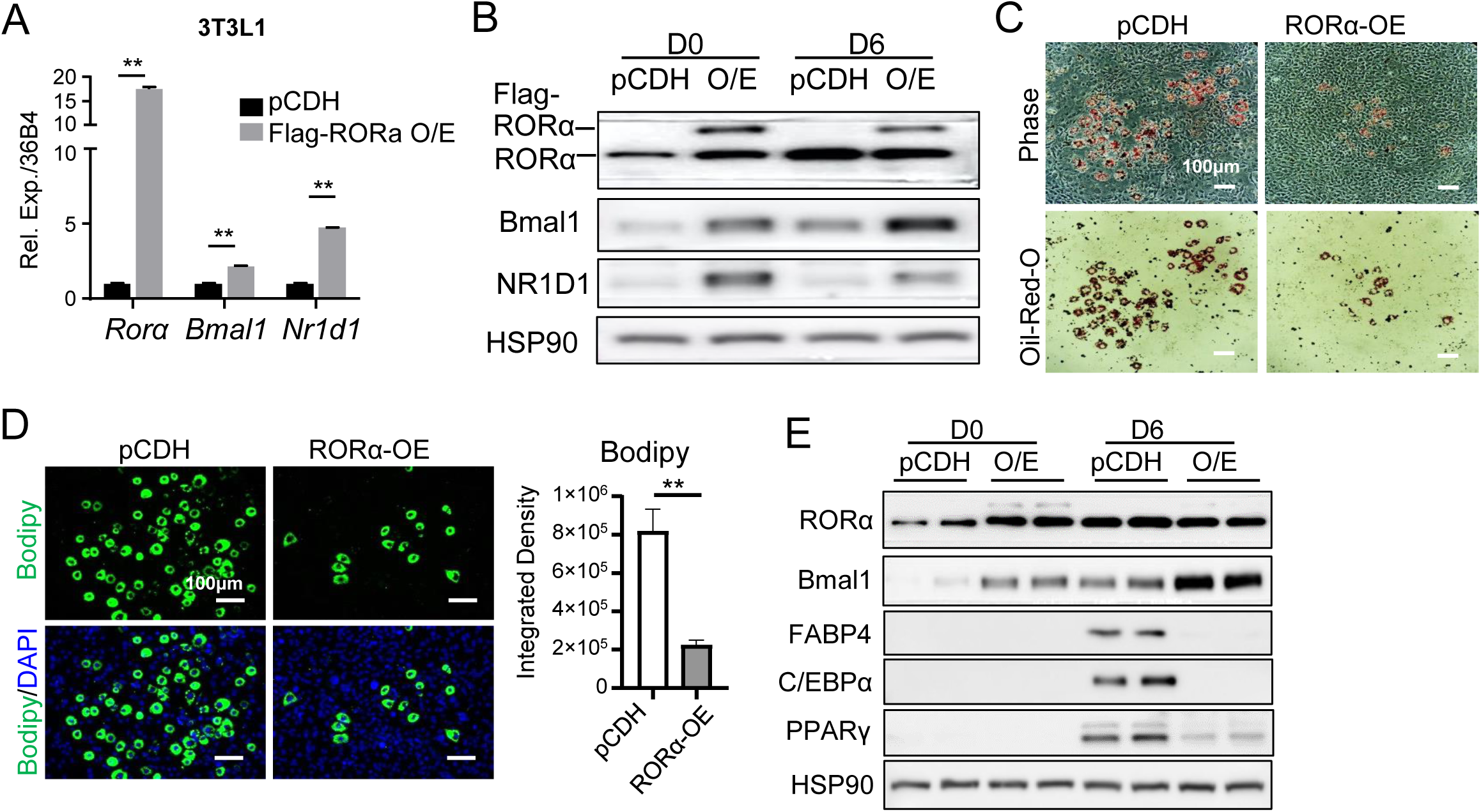
RORα gain-of-function in adipogenic progenitors inhibits adipogenesis. (A, B) Ectopic expression of RORα (RORα O/E) in 3T3-L1 preadipocytes induces clock gene activation as indicated by RT-qPCR (A) and immunoblot analysis (B). (C) Representative images of oil-red-O (C), and Bodipy staining with quantitative analysis (D), of adipogenic differentiation of 3T3-L1 preadipocytes containing pCDH vector control or RORα overexpression at day 6. Scale bar: 100 μm. (E) Representative images of immunoblot analysis of adipogenic protein expression of 3T3-L1 adipogenic differentiation with vector control or RORα overexpression following 6 days of adipogenic differentiation.

### In vivo efficacy of Quercetin and Fisetin on reducing fat mass and improving glucose metabolism

Based on the findings of quercetin and fisetin suppression of adipocyte development, we determined their potential impact on fat depots in vivo using wild-type C57/BL6 mice. Quercetin and fisetin were administered via intraperitoneal injections (50mg/Kg) for 5 consecutive days before analysis of body weight effects on distinct fat depots. Quercetin administration resulted in a slight reduction of body weight that did not reach statistical significance, potentially due to the short duration of the treatment, while fisetin administration had no effect on the body weight (Fig. 6A). Notably, the weight of both visceral and inguinal fat pads, representative of classic white adipose tissue and inducible thermogenic beige depots, respectively, were significantly reduced by quercetin, but were not affected by fisetin (Fig. 6B & 6C). In comparison, both compounds were able to attenuate the mass of interscapular brown fat (BAT, Fig. 5D), without affecting muscle weights as shown for tibialis anterior (TA, Fig. 6E) and gastrocnemius (GN, Fig, 6F) muscles. H/E histology of BAT from mice treated with Quercetin or Fisetin revealed marked reductions in lipid accumulation (Fig. 6G). Interestingly, both compounds displayed strong effects on reducing adipocyte hypertrophy of epidydimal white adipose tissue (eWAT, Fig. 6H) as well as inguinal beige fat pads (iWAT, Fig. 6I). In both fat depots, Quercetin displayed a more robust effect on attenuating adipocyte hypertrophy than that of fisetin, as indicated by the shift of size distribution toward smaller adipocytes.

**Figure 6.**
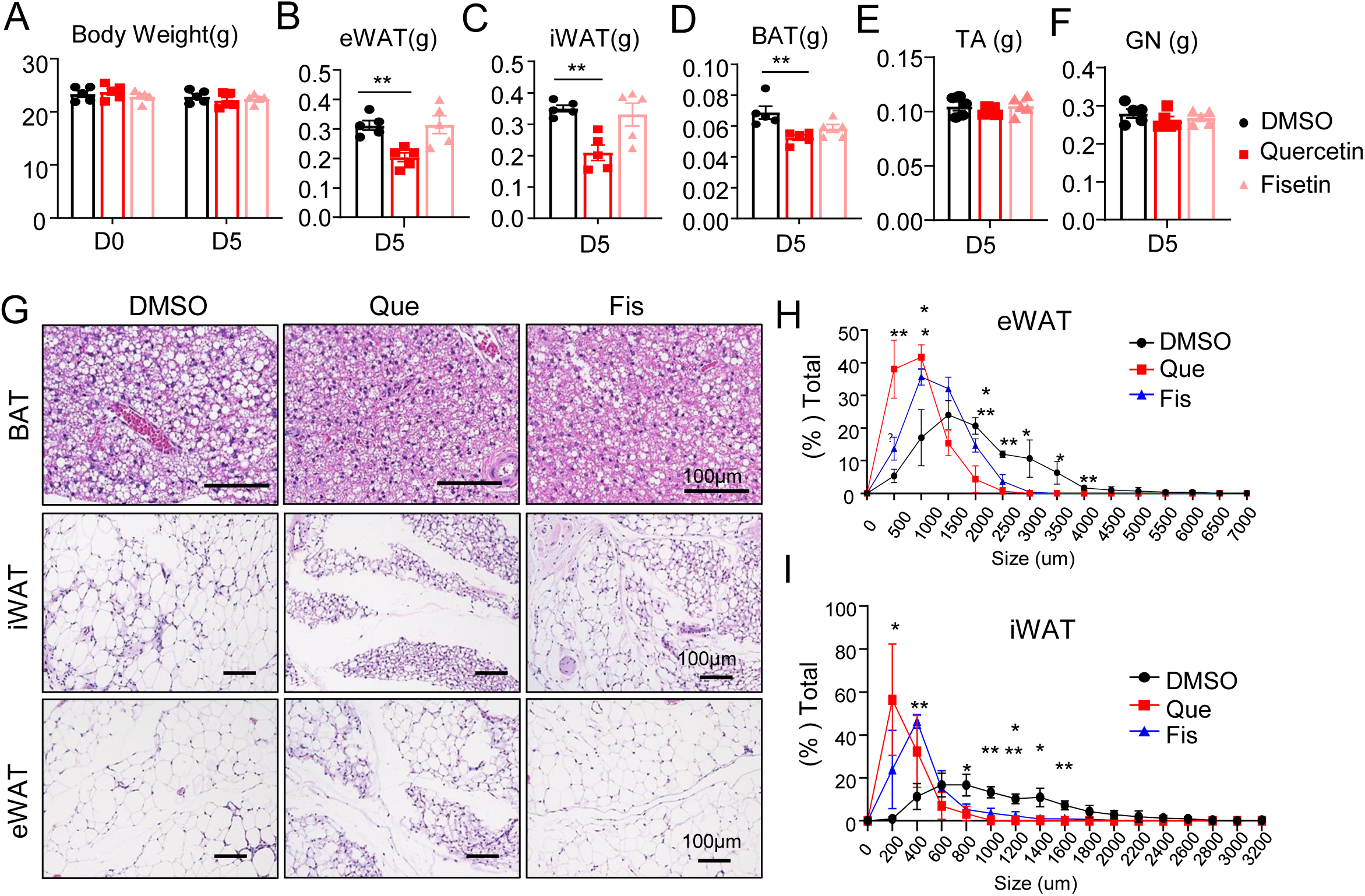
In vivo effects of Quercetin and fisetin treatment on reducing adipose tissue mass. (A-F) Effect of Quercetin and Fisetin on body weight (A) and individual tissue weight (B-F) in C57/BL6 mice via intraperitoneal delivery at 50mg/Kg for 5 days. (G-I) Representative H/E histology images of brown adipose tissue (BAT), inguinal and epidydimal white adipose tissue (iWAT & eWAT) in mice following 5 days of Quercetin or Fisetin treatment (G), with quantitative analysis of adipocyte cell size distribution of the eWAT (H) and iWAT (I). Scale bar: 100 μm. N=5/group. *, **: p<0.05 or 0.01 vs. DMSO by Student’s t test.

At the molecular level, we found that in both white (iWAT, Fig. 7A) and brown adipose depots (Fig. 7B), Quercetin and Fisetin were able to induce robust elevation of RORα protein expression along with Bmal1 induction by Fisetin in iWAT, suggesting their in vivo efficacy in activating clock function. To determine the underlying mechanisms mediating the in vivo effects of quercetin and fisetin, we analyzed the adipogenic, lipid remodeling and browning markers. Examination of iWAT samples revealed marked suppression of adipogenic factors (Fig. 7C) without altering PGC-1a level (Fig. 7D). In line with its more robust effect on reducing adipocyte hypertrophy, Quercetin displayed stronger inhibitions of adipogenic proteins than fisetin. In addition, there were inductions of the lipolytic enzyme adipose triglyceride lipase (ATGL) that were observed in this fat depot but not hormone-sensitive lipase (HSL, Fig. 7D), suggesting that enhanced lipid remodeling may contribute, at least in part, to the attenuated adipocyte hypertrophy in mice with quercetin or fisetin treatment. In BAT, both compounds markedly suppressed the adipogenic and lipogenic genes (Fig. 7 E), along with robust inductions of lipolytic enzymes ATGL and HSL as well as HSL phosphorylation (Fig. 7F). Notably, quercetin induction of lipolytic enzymes in BAT was also more robust than that of fisetin. Markers for brown fat thermogenic uncoupling activity, the Uncoupling Protein 1 (UCP-1) and the key factor in driving mitochondrial biogenesis, Peroxisome proliferator-activated receptor gamma coactivator-1α (PGC-1α) were not significantly altered by either quercetin or fisetin treatment (Fig. 7G). Analysis of serum metabolites revealed marked reduction of FFA levels by quercetin but not fisetin, suggesting improved systemic lipid metabolism in quercetin-treated cohorts (Fig. 7H), while circulating triglycerides were not affected by either treatment (Fig. 7I). Both quercetin and fisetin significantly reduced fasting plasma glucose to comparable levels (Fig. 7J). In addition, in response to insulin stimulation during the insulin tolerance test, quercetin-treated cohorts displayed significantly lower glucose concentration at 15 and 30 minutes, indicating improved insulin sensitivity (Fig. 7K).

**Figure 7.**
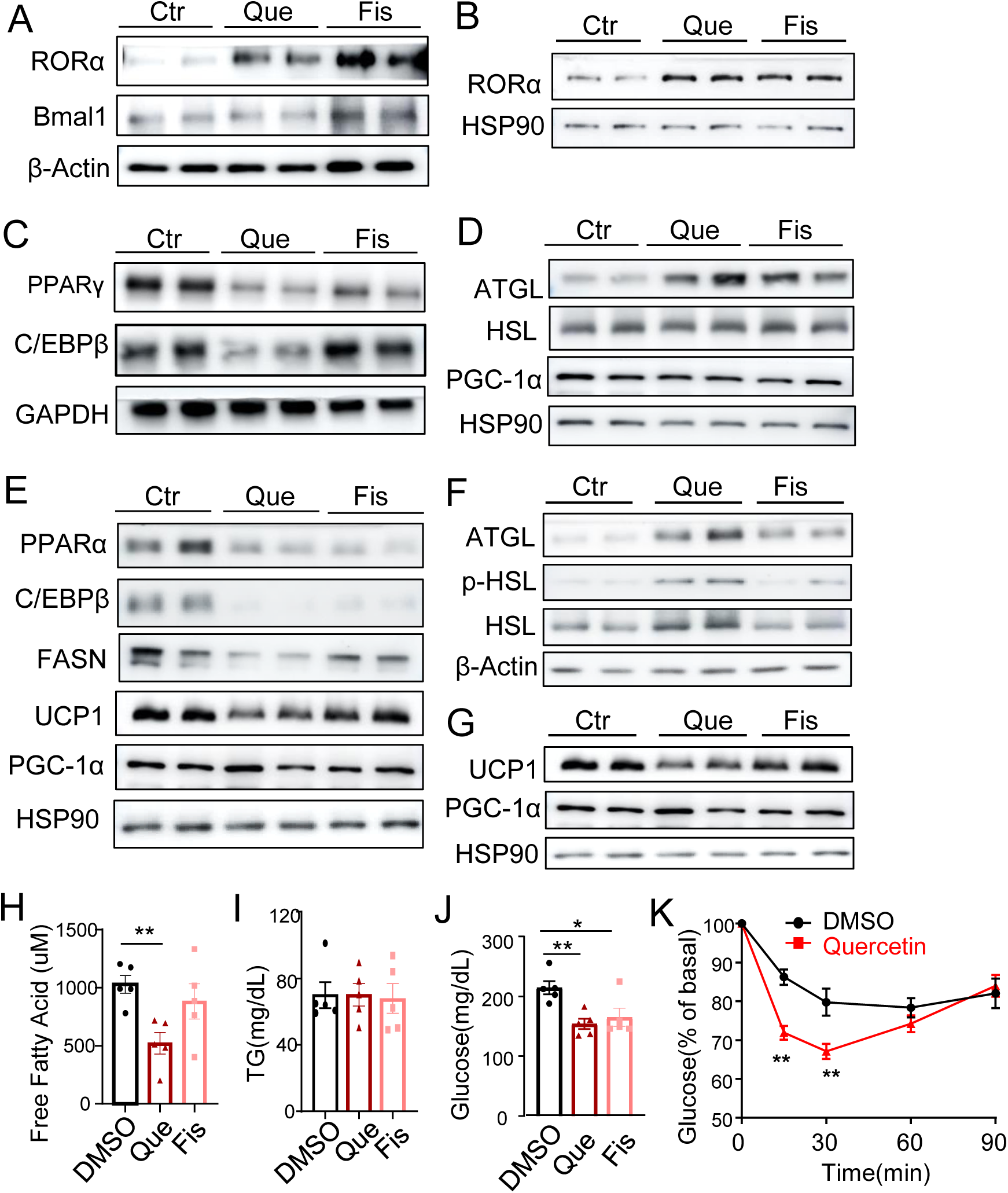
In vivo metabolic effects of Quercetin and Fisetin in adipose tissues. (A, B) Immunoblot analysis of Quercetin and Fisetin effect on inducing clock proteins in inguinal subcutaneous white adipose tissue (A) and brown adipose tissue (B). (C, D) Effect of Quercetin and Fisetin on adipogenic (C) and lipolytic proteins (D) in inguinal white adipose tissue of quercetin or fisetin-treated mice. (E-G) Representative immunoblot analysis of adipogenic (E), lipolytic (F) and thermogenic proteins (G) in BAT in quercetin or fisetin-treated mice. Each lane represents pooled sample of 2-3 mice/group. (H-J) Quercetin and Fisetin effect on serum free fatty acid (H), triglycerdie (I) and glucose (J) levels. (I) Insulin tolerance test as shown by glucose level expressed as percentage of basal value in control or Quercetin-treated mice following 0.75U/Kg of IP insulin injection. N=8/group. **: p< 0.01 vs. DMSO by one-way ANOVA with Tukey’s post-hoc test.

## Discussion

Dietary flavonoid compounds have established pleotropic metabolically beneficial effects, largely attributable to their anti-oxidant and anti-inflammatory properties [33, 49]. Our current study identified a novel mechanism of function of two structurally related flavonoids, quercetin and fisetin, as clock activators via RORα agonism, that directly suppress adipocyte development and attenuate lipid storage in mature adipocytes. Given the accumulating evidence that circadian clock dysfunction is risk factor contributing to the epidemics of metabolic disorders [3], re-enforcing clock via nutraceuticals may provide metabolic benefits to counteract the adverse consequence of circadian misalignments [7, 12].

Extensive studies to date have established the beneficial metabolic effects of flavonoid compounds. Our current finding of quercetin and fisetin as bone-fide RORα agonists provides additional mechanistic insights into their protective impact on metabolic regulations. We demonstrate that these molecules exert direct effects on inhibiting the adipogenic development of progenitor cells coupled with attenuating lipid accumulation in mature adipocytes. In vivo studies support these mechanisms with demonstrated effects on reducing fat tissue mass, with marked inhibition of adipogenic program and up-regulation of lipolytic pathways. Consistent findings on the blockade of adipogenesis by the synthetic ligand SR1078 and experiments using forced RORα expression further support the notion that RORα-mediated clock modulation in adipocytes may at least in part be responsible for the anti-obesity efficacy of quercetin and fisetin in vivo.

In differentiated adipocytes and in mice treated with quercetin, we found that quercetin was able to suppress lipogenic pathway while inducing lipolytic enzymes, suggesting that combined effects on limiting lipid synthesis while promoting mobilization from the adipocytes may underly the observed effect on reducing lipid storage and consequently hypertrophy. Given that adipose tissue hypertrophic expansion accounts for the major mechanism in obesity, these effects of quercetin in mature adipocytes could be a major mechanism responsible for the observed effects on reducing fat mass in vivo, while its inhibition of adipogenic and lipogenic drive in developing adipocytes may also contribute within the progenitor population during the development of obesity. Disruption of clock function, due to altered environmental lighting or a shiftwork schedule predispose to the development of obesity [20, 50–52] and are associated with a strong risk for insulin resistance and type II diabetes [4, 21, 52]. Discovery of these novel mechanisms of actions of quercetin and fisetin implicates their potential applications in circadian misalignment-associated disease conditions, such as people on shiftwork or the aging population that are prone to clock disruption-associated metabolic disorders [53, 54].

Circadian clock function is directly involved in modulating adipocyte biology. Bmal1 exerts direct transcriptional control of Wnt pathway that suppresses adipogenesis [17], while loss of clock function, due to ablation of either CLOCK or Bmal1 leads to the development of obesity and metabolic syndrome in mice [15–17]. Thus, maintaining or augmenting clock-controlled metabolic processes may offer new strategies to prevent or ameliorate obesity and associated metabolic consequences. Consistent with this idea, the related flavone compound nobiletin with RORα agonist activity [11] can suppress adipogenesis with demonstrated anti-obesity effect in mice [30]. Chlorhexidine, a newly-identified clock activator compound [55], also displayed robust anti-adipogenic effect along with its derivative [56]. By activating RORα, the clock-activating properties of quercetin and fisetin, may mediate, at least in part, their observed metabolic beneficial properties in vivo.

Quercetin and fisetin are known agents that promote cellular senescence, a class of molecules that are defined as senolytics [57]. The combination of Dasatinib and quercetin (D+Q) has been reported to show strong metabolic benefits in aged animals, with marked efficacy on reducing adiposity in aging cohorts [58]. Cellular senescence is an established marker of aging, and in aged adipocytes, senescence-associated secretory phenotype (SASP) can instigate the inflammatory milieu that exacerbates insulin resistance. The beneficial effects of Dasatinib and quercetin were largely attributed to their actions on eliminating senescent cells and the associated SASP consequence. Our findings of the clock-enhancing properties of quercetin and fisetin implicates this mechanism could be at play to mediate their metabolic effects, particularly in non-senescent cell populations within old tissues that comprise the majority of the tissue mass. Given that even in aged tissues/organs the population of senescent cells remain scarce, clock-modulatory benefits may contribute significantly to the overall metabolic effects of these compounds observed in vivo. Based on the strongest effects of quercetin and fisetin on reducing fat mass in vivo, we focused on their modulatory effects in adipocyte models, while their actions are likely not limited to these tissues. The effects mediated by clock or non-clock modulatory functions in other metabolic organs, such as the liver or skeletal muscle, could contribute to their overall systemic metabolic regulations in vivo.

Findings of quercetin and fisetin RORα agonist properties in modulating adipocyte behavior promoted our further investigations into RORα activation in adipocytes. Both pharmacological and genetic manipulation revealed inhibitory effects on adipogenic differentiation, consistent with the observed effects of quercetin and fisetin. RORα has an established role in blocking adipogenic differentiation via direct interaction and interference with the activity of adipogenic factor C/EBPβ [59]. Thus, quercetin and fisetin could function via blocking C/EBPβ activity to suppress adipocyte development, as supported by our in vitro and in vivo analyses. It is also possible that transcriptional effects of RORα on inhibiting lipid synthesis may contributes to effects in adipocytes. RORα suppresses the lipogenic program in hepatocytes, by blocking PPARγ activity that inhibits transcriptional inductions of SREBP-1c and Fasn [60], in line with observed inhibitory effects of FASN by quercetin and fisetin in adipose depots. Both quercetin and fisetin induced lipolytic enzyme expression in adipose tissue, suggesting that the breakdown of triglyceride storage in adipocyte may contribute to reduced adipocyte hypertrophy. Interestingly, circulating levels of free fatty acids were reduced by quercetin treatment with a tendency toward lower levels by fisetin. These findings together implicate additional effects of these compounds, potentially in other tissue sites that may promote the oxidation of fatty acids. AMPKα was reported to be activated by quercetin and fisetin that may also contribute to their anti-obesity effect [61], a potential mechanism that may underlie their beneficial metabolic in distinct metabolic organs.

Interestingly, the potency of quercetin and fisetin on activating RORα as shown by RORE-driven reporter are largely similar, only that the agonist activity of quercetin was maintained better at higher concentrations (5 and 10 µM) than that of fisetin. Their RORα-modulatory activities correlated with the observed in vivo efficacy, as quercetin effect on reducing adipocyte hypertrophy was more robust than that of fisetin. In contrast, their impacts on depleting lipid storage in brown fat were largely comparable. Analysis of adipogenic and lipogenic factors revealed stronger inhibition by quercetin than fisetin, whereas the induction of lipolytic enzymes displayed a similar tendency. Thus, the in vivo efficacy of quercetin as compared to fisetin, could be at least in part, mediated by their RORα agonist activity. However, as several types of oxysterols were identified as endogenous ligands of RORα [62, 63], it raises the possibility that quercetin and fisetin, depending on their effective intracellular concentrations, may compete with endogenous metabolite ligands for RORα binding dependent on cellular nutritional states. As the amount of flavonoid compounds, particularly quercetin and fisetin that are abundant in fruits and vegetables, are influenced by dietary intake, it is conceivable that supplementation in clinical applications may yield highly variable efficacies [64]. Discovery of clock activation by quercetin and fisetin with direct inhibitory effect on adipocyte development and hypertrophy adds to our current understanding of the metabolic benefits of these compounds.

The family of citrus flavonoids is extensive and includes the broad array of flavonols such as quercetin and fisetin, as well as largely apolar fully methoxylated flavones such as nobiletin and tangeretin [65]. Quercetin and fisetin displayed many functional similarities, but there were also differences worth noting. For example, the lipogenic inhibition of quercetin in cell culture models and its effects on body weight and gene expression modulation in vivo are more robust than those of fisetin. The structures of fisetin and quercetin differ by the presence of one hydroxyl group in the fifth cardon of the quercetin A ring. Thus, while some of the main biological effects of these compounds may be primarily related to the flavonoid backbone, certain effects could also be influenced by the type and number of functional groups present in these molecules [66]. Some of the differential effects noted between quercetin and fisetin underscore the importance of additional studies on the structural and functional relationships of individual flavonoids.

In summary, our study uncovered, for the first time, the anti-adipogenic effects of new clock-modulating compounds that may have potential for anti-obesity drug development. With the widespread metabolic consequences of circadian clock disruption, pharmacological targeting of the clock machinery to maintain metabolic homeostasis may offer new avenues for the prevention or treatment of obesity and related metabolic consequences.

## Acknowledgements

We thank Drs. Seung-Hee Yoo and Zheng Chen at University of Texas at Houston Health Science Center for providing the *Per2::dLuc* luciferase plasmids. We thank the City of Hope Shared Resources Animal Phenotyping for carrying out metabolic phenotyping analysis. KM is a faculty member supported by the NCI-designated Comprehensive Cancer Center at the City of Hope National Cancer Center. This project was supported in part by National Institute of Health grants R56AG080294 and an Arthur Riggs Diabetes and Metabolism Research Institute Innovative Award to KM. VY is supported by National Institute of Health grants R01DK097160 and R01DK128972, and AP is supported in part by National Institute of Health grant R01DK128972. These funders had no role in study design, data collection and analysis, decision to publish, or preparation of the manuscript.

## Data Availability Statement

All data generated in this study are included in this published article and associated supplementary information files. Reagents used in this study are available from the corresponding author upon reasonable request.

## Conflict of Interest

The authors declare that no competing interests exist that are relevant to the subject matter or materials included in this work.

## Ethics Statement

All procedures involving animal experiments were conducted in accordance with the protocol and ethical guidelines approved by the Institutional Animal Care & Use Committee (IACUC) of City of Hope protocol number: 17110.

## CRediT Author Contribution Statement

XX: methodology, data curation, investigation, formal analysis, validation, writing-review & editing; JP & TK: methodology, data curation, validation; KM: Conceptualization, visualization, formal analysis, validation, supervision, writing-original draft, review & editing, project administration, funding acquisition. ZS: resources, data curation, writing-review & editing; AP and VY: conceptualization, data curation, resources, writing-review & editing, funding acquisition.

## Notes

### Competing Interest Statement

The authors have declared no competing interest.

