## Supplemental tables for "Quercetin and Fisetin activate circadian clock via RORα and inhibit adipocyte growth"

**Supplemental Table 1. Primary antibodies list.**

| Antibody | Source | Cat# | Dilution |
|---|---|---|---|
| BMAL1 | Santa Cruz | SC-365645 | 1:1000 |
| CLOCK | Cell Signaling | 5157S | 1:1000 |
| RORα | Proteintech | 82930-1-RR | 1:1000 |
| DBP | Proteintech | 12662-1-AP | 1:1000 |
| NR1D1 | Proteintech | 14506-1-AP | 1:1000 |
| AMPKα | Cell Signaling | 2532S | 1:1000 |
| p-AMPKα(T172) | Cell Signaling | 2535S | 1:1000 |
| HSL | Cell Signaling | 4107S | 1:1000 |
| p-HSL(S563) | Cell Signaling | 4139S | 1:1000 |
| ATGL | Cell Signaling | 2439S | 1:1000 |
| UCP1 | Cell Signaling | 72298S | 1:1000 |
| PGC1α | Sigma | ST1204 | 1:1000 |
| C/EBPα | Santa Cruz | SC-61 | 1:1000 |
| C/EBPβ | Santa Cruz | SC-7962 | 1:1000 |
| PPARγ | Santa Cruz | SC-7273 | 1:1000 |
| FASN | Santa Cruz | SC-48357 | 1:1000 |
| FABP4 | Santa Cruz | SC-271529 | 1:1000 |
| GAPDH | Proteintech | 60004-1-IG | 1:3000 |
| HSP90 | Cell Signaling | 4874S | 1:3000 |
| β-Actin | Proteintech | 66009-1-Ig | 1:3000 |

**Supplemental Table 2. Primer sequence for qPCR analysis.**

| Genes | | Sequences |
|---|---|---|
| Rorα | Forward | GAACACCTTGCCCAGAACAT |
| | Reverse | TTGGCAAACTCCACCACATA |
| Bmal1 | Forward | CGCTTTCTGGAGGGTGTCCGC |
| | Reverse | TGCCAGGACGCGCTTGTACC |
| Nr1d1 | Forward | TGGCATCCGGTGCACTGCAG |
| | Reverse | CCCTCCAGAAGGGTAGCACGCT |
| 36B4 | Forward | CGCTTTCTGGAGGGTGTCCGC |
| | Reverse | TGCCAGGACGCGCTTGTACC |
